# When the Good Guy Becomes the Bad Boy: Social Information Modulates the Neural, Physiological and Subjective Responses to Emotional Facial Expressions

**DOI:** 10.1101/077834

**Authors:** Martial Mermillod, Delphine Grynberg, Magdalena Rychlowska, Nicolas Vermeulen, Paula M. Niedenthal, Brice Beffara, Léo Lopez, Sylvie Droit-Volet

## Abstract

In the past decade, different studies have suggested that high-order factors could influence the perceptual processing of emotional stimuli. In this study, we aimed to evaluate the effect of congruent vs. incongruent social information (positive, negative or no information related to the character of the target) on subjective (perceived and felt valence and arousal), physiological (facial mimicry) as well as on neural (P100 and N170) responses to dynamic emotoional facial expressions (EFE) that varied from neutral to one of the six basic emotions. Across three studies, the results showed (1) reduced valence and arousal evaluation of EFE when associated with incongruent social information (Study 1), (2) increased electromyographical responses (Study 2) and significant modulation of P100 and N170 components (Study 3) when EFE were associated with social (positive and negative) information (vs. no information). These studies revealed that positive or negative social information reduced subjective responses to incongruent EFE and produces a similar neural and physiological boost of the early perceptual processing of EFE irrespective of their congruency. In conclusion, this study suggested that social context (positive or negative) enhances the necessity to be alert to any subsequent cues.

## Introduction

Behavioral evidence suggests the relevance of top-down processes in the use of concepts (Schyns, Goldstone, & Thibaut, 1998), percepts (Bar, 2004; Vermeulen, Mermillod, Godefroid, & Corneille, 2009) and affects (Hess, Adams, & Kleck, 2007; Niedenthal, Winkielman, Mondillon, & Vermeulen, 2009; Rudrauf, David, Lachaux, Kovach, Martinerie, Renault, & Damasio, 2008; Scherer, 1997). For instance, participants’ personality (e.g., Campanella, Falbo, Rossignol, Grynberg, Balconi, Verbanck, & Maurage, 2012) or facial expressions (Niedenthal, Winkielman, Mondillon, & Vermeulen, 2009) modulate the detection or recognition of emotional facial expressions (EFE). In addition, research has pointed to a modulation of EFE processing by social factors, such as the degree of friendship between the participant and the target (Hess, et al., 1995) or the social norms of the participants (Scherer, 1997).

In a social neuroscience perspective, recent studies show that the physiological (i.e., facial mimicry) and neural underpinnings of the perception of EFE may also depend upon high-level social factors associated with the targets or with the participants. For instance, facial emotions expressed by cultural in-group (compared to out-group) targets modulate facial mimicry of participants (e.g., Mondillon, Niedenthal, Gil, & Droit-Volet, 2007) and event-related potentials in response to these EFE (Kubota & Ito, 2007). In another study, happy expressions of targets presented as future interaction partners (socially relevant) lead to enhanced late positive potentials (LPP) amplitudes relative to non socially relevant targets (Bublatzky, Gerdes, White, Riemer, & Alpers, 2014). Because LPP amplitudes are larger for high arousing stimuli (vs low arousing) (Schupp, Flaisch, Stockburger, & Junghöfer, 2006), these results suggest that social relevance may act as top-down factor on positive EFE, leading observers to perceive happy faces as more relevant (i.e., larger LPP amplitude).

Together, these studies suggest that social factors associated with the participant, the target or with the social interaction between them may influence the processing of EFE at subjective (recognition accuracy), physiological (facial mimicry) and neural (event-related potentials) levels. However, most of these studies have either focused on the participants’ characteristics (e.g., Likowski et al., 2008) or manipulated the social appraisal of the targets by varying perceptual features (e.g., ethnicity) of the face itself (e.g., Kubota & Ito, 2007) or the nature of the interaction between the participant and the target (e.g., Bublatzky et al., 2014). There is so far few evidence that the subjective, physiological and neural responses to EFE are influenced by high-order social information about the person depicted in the EFE. To our knowledge, only one study investigated whether positive, negative or neutral characteristics of the target modulated facial mimicry of EFE (Likowski, Mühlberger, Seibt, Pauli & Weyers, 2008). This study revealed that positive traits (e.g., kind) enhanced Zygomatic activity for happy expressions and Corrugator activity for sad expressions. Moreover, negative (e.g., malicious) and neutral (e.g., neutral) traits had no influence on Zygomatic traits but reduced Corrugator activity for sad expressions. Although this study highlight the influence of positive or negative traits on EMG responses to EFE (i.e;, enhanced mimicry for positive targets), each avatar had their specific traits thus preventing the control for the perceptual stability of facial expressions.

Therefore, the aim of the present study was to examine whether high-level social information about the target (1) influences subjective responses to EFE, (2) impacts quick mimicry and perceptual processes implied in EFE processing, and (3) by keeping stable the perceptual features. The investigation of such influence is highly relevant as recent models propose that the psychological and neural underpinnings of the perception of EFE depend upon the understanding of social context (Niedenthal et al., 2010). Furthermore, we postulate that the effect of social information on EFE processing could be interpreted within the theoretical framework of emotional embodied simulation (Barsalou, 1999; Dimberg, 1990; Hess, Banse, & Kappas, 1995; Niedenthal, 2007), which assumes that the processing of emotional information is grounded in the brain’s perceptual, affective and sensory-motor systems. Several studies have shown that the processing of emotional concepts (e.g., emotional words or EFE) leads individuals to simulate this concept at a bodily level (e.g., Niedenthal, Winkielman, Mondillon, & Vermeulen, 2009). Indeed, preventing participants from simulating emotional concepts (i.e., blocking participants’ facial expressions) during EFE recognition task reduced recognition accuracy (Havas, Glenberg, Gutowski, Lucarelli, & Davidson, 2010). Finally, it has been shown that the recognition of emotional facial expressions is influenced by the activation (or inhibition) of facial muscles used to express the perceived emotion (Niedenthal, 2007). Similar findings were reported with direct TMS modulation of the sensorimotor cortex (Pitcher, Garrido, Walsh, & Duchaine, 2008). Together, these studies support that EFE are bodily grounded and that their processing is modulated through the embodied simulation. In respect to the moderating role of social information on the effect of EFE on subjective, physiological and neural responses, we postulate that through their influence on embodied simulation of EFE, social factors associated with the target may influence their processing.

More precisely, the aim of the present research was to examine whether social information as well as its valence (positive vs negative) modifies the evaluation of the valence and arousal of EFE, as well as facial imitation and early neural components related to perceptual information (P100 and N170). This is based on previous findings that mimicry processes assessed by electromyography (EMG) activity and P100 and N170 amplitudes were modulated by the expressed emotion during a EFE recognition task (Achaibou, Pourtois, Schwartz and Vuilleumier, 2008). Therefore, the present study aimed to replicate these results and to investigate the top-down influence of social information during perception of EFE on their subjective, physiological and neural responses. The P100 component, occurring approximately 120 msec after the stimulus onset, is associated with the processing of exogenous visual stimulation in extrastriate cortical areas (Clark & Hillyard, 1996), suggesting that the P100 is modulated by the physical properties of the stimulus. In respect to EFE processing, the exogenous nature of the P100 component has been supported by previous studies showing that fearful emotions generated larger amplitudes as compared to neutral emotions (Pourtois, Dan, Grandjean, Sander, Vuilleumier, 2005; Pourtois, Grandjean, Sander, & Vuilleumier, 2004). However, the emotional stimuli used in these studies were different at the emotional (endogenous) and at the perceptual (exogenous) level. For instance, angry vs. happy faces differ at the emotional level, but they are also considerably different at a purely perceptual level, even after a careful control of luminance or contrast information. Therefore, the modulation of the P100, as well as the modulation of N170 component (related to facial processing) by endogenous variables, (e.g., social information) remains a matter of debate. Thus, in addition, the current study also aimed to determine whether the P100 and N170 components could be modulated by endogenous social factors applied on *strictly identical EFE at a perceptual level.* Furthermore, at a physiological level, we aimed to evaluate whether mimicry processes, assessed by EMG responses to EFE, is modulated by social information.

## Hypotheses

**Study 1** examined the effect of a positive, negative or no social information on participants’ ratings of the valence and intensity of EFE. We expected significantly higher ratings of valence and intensity when the valence of the EFE was congruent with the valence of the social label (e.g. joy expressed by nice nurse) compared to the control situation without social information. Conversely, we expected lower ratings of valence and intensity for a given EFE when the valence of the EFE and the social label (i.e. joy expressed by a serial rapist) were incongruent compared to the control situation without social label. We will also examine how the emotion *felt* by the participants is modulated by social information. Indeed, it has been shown that the social information (i.e., ethnicity) about the target depicted in the EFE modulates empathic emotional responses (Drwecki et al., 2011).

**Study 2** investigated the impact of social information on EMG activity. We recorded EMG over six facial muscles involved in specific expressions, such as the contraction of the zygomaticus major (i.e., in response to happiness) and the corrugator supercilii (i.e., in response to anger). We expected increased EMG activity of the corresponding muscle when the facial expression was preceded by social information congruent to the emotion displayed by the stimulus compared to no social label. For example, we expected that congruency trials (i.e., happy facial expression with a positive label) should elicit higher EMG activity in muscles involved with producing facial expressions of joy (zygomaticus major and orbicularis oculi) compared to the same EFE presented without social information. This hypothesis is based on previous studies showing that social factors may modulate facial mimicry (Likowski, Mühlberger, Seibt, Pauli & Weyers, 2007; Mondillon et al., 2007). Conversely, incongruent trials (e.g., happy facial expression with a negative label) should produce lower EMG activity of the corresponding facial muscles (zygomaticus major and orbicularis oculi) compared to the same EFE with no social information.

**Study 3** examined the potential effects of social information on two early visual ERP components, namely the P100 and N170. Based on Achaibou et al. (2008), we expected higher amplitudes of the P100 and N170 when the EFE is congruent with the preceding social information compared to the control situation in which no social information is provided. Conversely, incongruent trials (e.g., happy facial expression with a negative label) should produce lower amplitudes of the P100 and N170 compared to the same EFE with no social information.

### Study 1

The influence of social context on the processing of emotional faces was first investigated at the subjective level. The goal was to examine the effects of social context on the ratings of valence and intensity of the EFE as well as the emotions felt in response to these EFE.

## Method

### Participants

Twenty-four undergraduate students (19 female) (*M*_*age*_ = 20.50, *SD*_*age*_ = 2.54) at the Blaise Pascal University, Clermont-Ferrand (France) with corrected-to-normal vision, participated in exchange for course credits. All participants gave written informed consent and had normal or corrected vision and no psychiatric or neurological disorders.

### Material and Procedure

<u>*Social information*.</u> The social context of EFE was operationalized with verbal labels referring to positive and negative categories. Each label was composed of two words (noun and adjective; e.g., loving mother). Thirty-two labels were initially created: 16 positive and 16 negative adjectives referring to either male or female nouns (see Appendix). A pilot study was carried out with 24 undergraduate psychology students (20 females; *M*_*age*_ 20.70; *SD*_*age*_ = 2.54) to test to pleasantness and unpleasantness of all labels. For each label, participants rated the extent to which the person described by the label was pleasant or unpleasant with two 7-point Likert scales for pleasantness and unpleasantness (ranging from 0 to 6; not at all to totally). The six labels rated as most pleasant (i.e., loving mother, passionate teacher, humanitarian doctor, nice nurse, cheerful sportsman, caring father; *pleasant scale*: *M* = 5.76, *SD* = 0.29*; unpleasant scale: M* = 2.06, *SD* = 0.37) and six rated as most unpleasant (i.e., dangerous schizophrenic, sadistic killer, serial rapist, asocial necrophiliac, brutal trainer, paedophile therapist; *pleasant scale*: *M* = 1.63, *SD* = 0.49*; unpleasant scale: M* = 6.30, *SD* = 0.58) were retained for the experiments. Subsequent analysis showed that positive labels significantly differed from negative labels in terms of pleasantness (*t*(10) = 17.69, *p* <.001) and unpleasantness (*t*(10) = −15.12, *p* < .001).

<u>*Dynamic EFE*</u>. The dynamic faces expressed a neutral emotion and then either gradually changed to express the full intensity of 6 basic emotions (anger, disgust, fearful, happiness, sadness and surprise; Ekman & Friesen, 1976) or remained neutral (no emotion). For each emotional category, there were ten different face identities (4 females and 6 males). This made a total of 70 different movie clips (10 identities x 7 EFE). For a reliable control of the onset, duration, and intensity of emotional expressions, we used dynamic expressions from a set of morphed pictures from a neutral to an emotional expression of the same face identity on the basis of Benson and Perrett’s morphing technique (Benson & Perrett, 1993). Thus, we used 18 frames per face with increasing emotional intensity for each emotion. Each of the first 17 frames was exposed for 40 msec and the 18^th^ frame lasted for 2000 msec (Figure 1).

**Figure 1.**
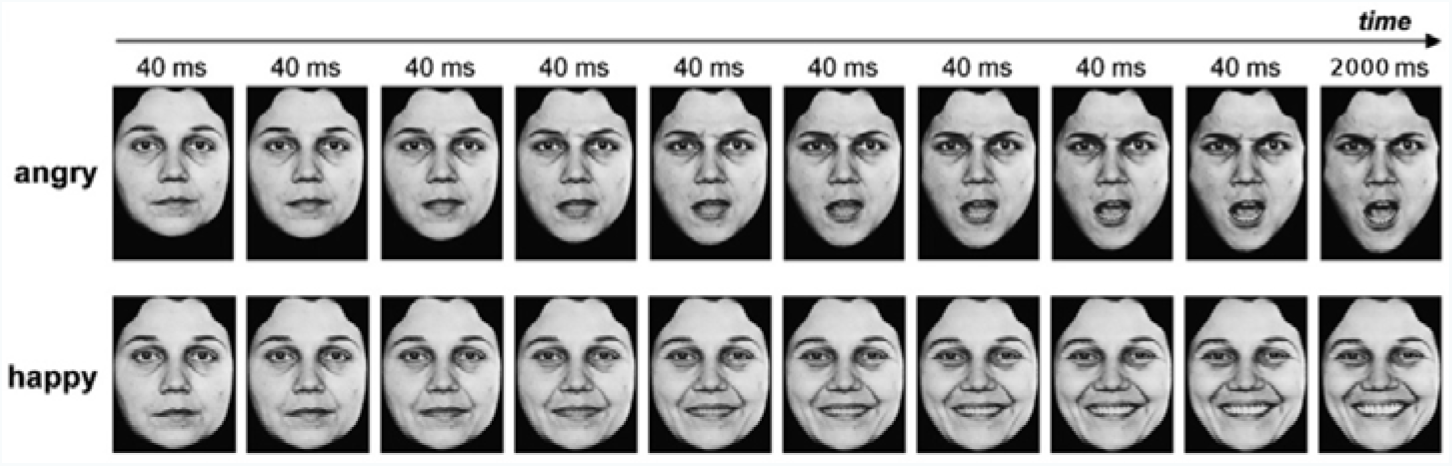
Example of dynamic facial expressions.

### Procedure

Participants were seated in front of a computer screen on which the dynamic EFE was displayed in central vision. EFE pictures were presented using the E-Prime software (Psychology Software Tools, http://www.pstnet.com) creating the compelling illusion of a short movie clip displaying a dynamic facial expression from neutral to emotional expressions. The experiment was divided into two phases. In the first experimental phase, participants were exposed to a central fixation cross for a duration varying from 500 msec to 1000 msec, followed by a video presenting one EFE (2680 msec). After each of the seventy videos, participants were instructed to rate the EFE in terms of valence on a continuous pixel scale ranging from 0 (positive) to 100 (negative). Next, participants rated the intensity of the EFE on each of the six basic emotions (anger, disgust, fear, happiness, sad, or surprise) on an identical pixel scale (ranging 0: not at all, to 100: totally).The second phase was similar to the first phase, except each of the seventy videos was preceded by a positive or negative label. A specific identity from the database (for every EFE expressed by this individual) was associated with a positive label (e.g., nice nurse) for half of the participants, and a negative label for the other half (e.g., serial rapist). The social valence of each identity was counterbalanced across participants. After seeing the label displayed on the screen for 2000 msec, participants were exposed to a fixation cross (from 500 msec to 1000 msec) before each video depicting a specific EFE of a specific person (for 2680 msec). Similarly to phase 1, participants had to assess the valence and the intensity of the *perceived* EFE. Moreover, participants were also instructed to evaluate the emotion they *felt* after exposure to each video on the same continuous scales as Phase 1 (i.e., valence and intensity). The presentation order of movie clips was randomized for each phase.

### Statistical analyses

All analyses were performed using PASW Statistics 18 (SPSS Inc., Chicago, IL). We conducted an analysis of variance (ANOVA) on valence and intensity of the emotion *perceived* (Phase 1 and 2) and valence and intensity of the emotion *felt* by the participant (for Phase 2 only) as the dependent variables, and EFE (Anger; Disgust; Fear; Happiness; Sad; Surprise or Neutral) and social label (Phase 1 - no social label; Phase 2 - negative social label; Phase 2 - positive social label) as within-subjects independent variables. Concerning the emotional intensity of EFE, for simplicity and clarity reasons, we only kept the intensity level related to each emotion (e.g., anger responses after presentation of an angry faces) and not the entire confusion matrices.

### Perceived Valence

At the level of valence, we observed a significant two-way interaction, *F*(10, 230) = 5.38, *p*<0.001, between social label and EFE (Table 1). Pairwise comparisons revealed that facial expressions of anger, *F*(1, 23) = 21.85, *p*<.001, disgust, *F*(1, 23) = 23.09, *p*<.001, fear, *F*(1, 23) = 12.04, *p*<.01 and sadness, *F*(1, 23) = 13.83, *p*<.01, were evaluated as less negative when preceded by positive social label than without social label. Conversely, facial expressions of happiness, *F*(1, 23) = 11.05, *p*<.01, and sadness, *F*(1, 23) = 10, *p*<.01, were evaluated as less positive when preceded by negative social label than without social label. Other pairwise comparisons were not significant.

**Table 1.** Average rated level of perceived valence, from 0 (negative) to 100 (positive), of each emotional expression depending on the valence of the label (negative, positive or no label).

| Displayed<br>EFE | Social label |  |  |
| --- | --- | --- | --- |
|  | No social<br>valence | Negative social<br>valence | Positive social<br>valence |
| Anger | 9.19 | 10.96 | 16.77 |
| Disgust | 11.08 | 12.15 | 20.05 |
| Fear | 12.26 | 14.9 | 18.55 |
| Happy | 92.46 | 81.4 | 90.24 |
| Sad | 11.92 | 17.8 | 18.26 |
| Surprise | 43.16 | 40.63 | 45.59 |

### Perceived Emotional Intensity

Analyses of the intensity of the perceived emotional expressions revealed a significant two-way interaction, *F*(10, 230) = 2.01, *p*<.05, between social label and EFE (Table 2) on the intensity of the corresponding emotion. Pairwise comparisons revealed that, compared to no label, a negative label significantly decreased the perceived intensity for disgust, *F*(1, 23) = 6.73, *p*<.05, surprise, *F*(1, 23) = 4.59, *p*<0.05, and marginally for happiness, *F*(1, 23) = 3.52, *p*=.07. Conversely, disgust, *F*(1, 23) = 22.28, *p*<.001 and fear, *F*(1, 23) = 8.8, *p*<.01, were rated as less intense when preceded by positive social label than without social label. The effect was marginally significant for the expression of anger, *F*(1, 23) = 3.28, *p*=.08. Other pairwise comparisons were not significant.

**Table 2.** Average ratings of intensity of *perceived* emotional expression, from 0 (not expressed) to 100 (fully expressed), as a function of the valence of labels (negative, positive or no labels).

| Displayed<br>EFE | Social label |  |  |
| --- | --- | --- | --- |
|  | No social label | Negative social label | Positive social label |
| Anger | 69.43 | 70 | 63.59 |
| Disgust | 57.27 | 45.22 | 43.09 |
| Fear | 59.1 | 54.7 | 50.29 |
| Happy | 87.22 | 81.47 | 86.4 |
| Sad | 60.25 | 59.35 | 56.85 |
| Surprise | 83.4 | 76.24 | 77.59 |

### Felt valence

We observed that the social valence of the stimulus affected the participants’ responses concerning felt valence. More precisely, results point to a significant interaction between the social information and EFE (*F*(5, 115) = 20.6, *p*<0.001). To describe this interaction effect, we performed pairwise comparisons, which revealed that negative labels were associated with more negative feelings than were positive labels for anger (*M*_*negative* *social* *label*_= 18.38, *SE* = 3.14 vs. *M_positive social label_*= 32.49, *SE* = 3.31), *F*(1, 23) = 23.86, *p*<0.001, disgust (*M_negative social label_*= 17.82, *SE* = 2.33 vs. *M_positive social label_*= 35.48, *SE* = 3.28), *F*(1, 23) = 27.89, *p*<0.001), and surprise (*M*_*negative* *social* *label*_= 37.68, *SE* = 2.95 vs. *M*_*positive* *social* *label*_= 50.56, *SE* = 3.02), *F*(1, 23) = 16.32, *p*<0.001). Positive labels were associated with more positive feelings than were negative labels for happiness (*M*_*negative* *social* *label*_= 44.75, *SE* = 4.48 vs. *M*_*positive* *social* *label*_= 81.04, *SE* = 3.09), *F*(1, 23) = 55.43, *p*<0.001). No significant differences were observed for facial expressions of fear and sadness.

### Felt Emotional Intensity

We also found a significant interaction between social label and EFE (*F*(5, 115) = 17.80, *p*<0.001) regarding the intensity of corresponding emotion felt by the participants. Participants felt greater anger after negative relative to positive labels (*M*_*negative* *social* *label*_= 22.77, *SE* = 5.44 vs. *M*_*positive* *social* *label*_= 11.95, *SE* = 4.74), *F*(1, 23) = 11.13, *p*<0.01. We found a similar effect for disgust (*M*_*negative* *social* *label*_= 31.74, *SE* = 5.72 vs *M*_*positive* social label_= 17.17, *SE* = 4.81), *F*(1, 23) = 18.48, *p*<0.001) and sadness (*M*_*negative* *social* *label*_= 19.1, *SE* = 4.9 vs *M*_*positive* *social label*_= 31.72, *SE* = 4.79), *F*(1, 23) = 13.3, *p*<0.01). Conversely, participants reported a higher level of happiness after positive labels compared to negative labels (*M*_*negative* *social* *label*_= 19.1, *SE* = 4.9 vs. *M*_*positive* *social* *label*_= 53.72, *SE* = 6.31), *F*(1, 23) = 26.63, *p*<0.001). There was not a significant difference between positive and negative social label for fear and surprise emotions.

### Discussion

In respect to *perceived valence* of facial expressions, this study showed that expressions of anger, disgust, fear and sadness were perceived as less negative when preceded by a positive label compared to no label. Conversely, facial expressions of happiness as well as sadness were rated as less positive when preceded by a negative label compared to no label. Regarding the ratings of the *intensity perceived of the emotional expressions,* expressions of disgust, surprise and happiness were rated as less intense when preceded by negative social labels compared to no label. Expressions of disgust, fear and anger were also rated as less intense when preceded by positive social labels.

Concerning ratings of the *valence of the emotion felt*, we showed similar results. Participants reported less negative valence for anger, disgust and surprise and more positive valence for happiness emotions expressed by an individual associated with positive (compared to negative) social information. In their ratings of the *felt emotional intensity*, participants reported higher feelings of anger, disgust, sadness, for individuals presented with the negative social label compared to positive social label. Conversely, participants reported higher levels of happiness in response to individuals associated with a positive label compared to a negative label.

This study thus supports hypothesis 1 according to which high-order target-related social information influences the ratings of valence and intensity of the EFE as well as the emotions felt in response to these EFE. More precisely, incongruent social terms reduce the evaluation of valence and intensity of the EFE and congruent trials led to higher intensity of felt emotions. Moreover, incongruent trials led to a lower valence of felt emotions compared to a control situation of perceiving emotional expressions without social information. We thus extend upon previous findings suggesting that top down social factors influence EFE processing (e.g., Bublatzky et al., 2014; Kubota & Ito, 2007; Likowski et al., 2008). In terms of affective responses to EFE, we support Drwecki et al.’s (2011) findings that emotional responses are influenced by social information (i.e., ethnicity) about the target depicted in the EFE.

More globally, this study is in line with study showing the influence of participants’ appraisal of the target on the emotional evaluation and responses to EFE. For instance, Lamm and colleagues (2007) assessed how reappraisal might influence the participants’ evaluation of the pain expressed by faces of patients undergoing painful treatment. When participants were told that the treatment was effective (versus non effective), they judged the experience of the person as being less painful. Results also showed that participants from the "non effective" group reported higher distress than those from "effective" group. Therefore, in line with Lamm et al’s (2007) results, the present findings showed for the first time that appraising EFE as socially congruent or incongruent with the expression modulates their evaluation as well as their emotional effects.

Study 2 and 3 aim to determine the neural and physiological processes underpinning the modulation of social information of EFE on responses to these EFE. According to the embodiment theory, we assume that the embodiment processing of EFE might be modulated by social information, such that participants might have less simulated EFE associated with incongruently social information.

### Study 2

In this study, we investigated the impact of social information on mimicry within the theoretical framework of the embodiment theory. More specifically, we expected increased EMG activity in the corresponding muscles when EFE was preceded by a congruent social label compared to an EFE without a social label. Conversely, incongruent trials should produce lower EMG activity of the corresponding facial muscles compared to the same EFE with no social label.

## Method

### Participants

Twenty healthy subjects (*M*_*age*_ = 20.80 years, *SD*_*age*_ = 2.26) at the Blaise Pascal University, Clermont-Ferrand (France), with corrected-to-normal vision, participated in exchange for course credits. All participants gave written informed consent and had no psychiatric or neurological disorders.

### Material and procedure

The experimental design of Study 2 is similar to study 1, except that we recorded participants’ EMG activity during the experimental task. Electrodes were fixed on the participant’s face in order to record facial muscle electrical activity (EMG). Before attaching the electrodes, the skin was cleaned with alcohol in order improve the signal (i.e, removing sebum). Six pairs of electrodes were placed on the following muscle regions (see Fridlund & Cacioppo, 1986 for details): frontalis pars medialis (mainly related to surprise and fear), corrugator supercilii (mainly related to anger), orbicularis occuli pars orbitalis (mainly related to happiness), levator labii superioris (mainly related to disgust), zygomaticus major (mainly related to smile) and orbicularis oris inferior (mainly related to sadness). The reference electrode was clipped on the ear lobe. Because of the large electrode size, both sides of the face were required. Half of the electrodes (corrugator, levator and orbicularis oris) were placed on the right side of the face while the other three electrodes (frontalis, orbicularis occuli, zygomaticus major) were placed on the left side. After checking the position of each pair, study instructions were given to the participants and the experiment began.

### Data acquisition

Each EMG signal was acquired by bipolar electrodes used for electrophysiological acquisition and amplified by Multi-Channel Bio Amps GT201 band pass filtered from 10 to 1000 Hz (ADInstruments equipment, ML880 Powerlab 16/30). The EMG data were averaged during the 5000 msec after stimulus onset. The dependent variable of interest was measured as the difference between the mean activity 5000 msec after the stimulus and the baseline activity recorded during the 500 msec before the stimulus onset, when a fixation cross was displayed on the screen.

### Statistical analyses

All analyses were performed using PASW Statistics 18 (SPSS Inc., Chicago, IL). We conducted an analysis of variance (ANOVA) with EMG activity (6 levels: frontalis pars medialis, corrugator supercilii, orbicularis occuli pars orbitalis, levator labii superioris, zygomaticus major and orbicularis oris inferior) as dependent variables and EFE displayed on screen (Anger; Disgust; Fear; Happiness; Sad; Surprise or Neutral) and social label (no social label; negative social label; positive social label) as within-subjects independent variables. Corrections of Greenhouse-Geisser were also applied for variance analysis when violations of sphericity occurred.

## Results

We observed a significant interaction between emotion and muscle, *F*(5.54, 105.22) = 2.37, *p*< .05, confirming, as reported in preliminary analysis (Beffara et al., 2012), that each emotion was related to a specific set of muscles (e.g. zygomaticus major with joy, corrugator supercilii with anger). Therefore, we focused the analyses on the muscles specific to each emotion (Figure 2). Statistical analyses revealed a significant main effect of social label, *F*(1.36, 25.36) = 8.32, *p*< .01. As shown on Figure 2, labels of positive and negative social valence produced a significant increase in EMG activity in response to EFE compared to the EMG activity elicited by EFE displayed to the control situation without social information. Negative (*t*(19) = 3.34, *p*< .01, two-tailed t-test) and positive (*t*(19) = 2.83, *p*< .05, two-tailed t-test) social labels produced a significant increase in EMG activity compared to control situation without social labels while the effect of positive and negative social labels did not differ (*t*(19) = 0.80, *p*=.93, two-tailed t-test). No main effect was observed for either emotion or muscle. All the remaining interactions (emotion x social label; muscle x social label; muscle x emotion x social label) were not significant.

**Figure 2.**
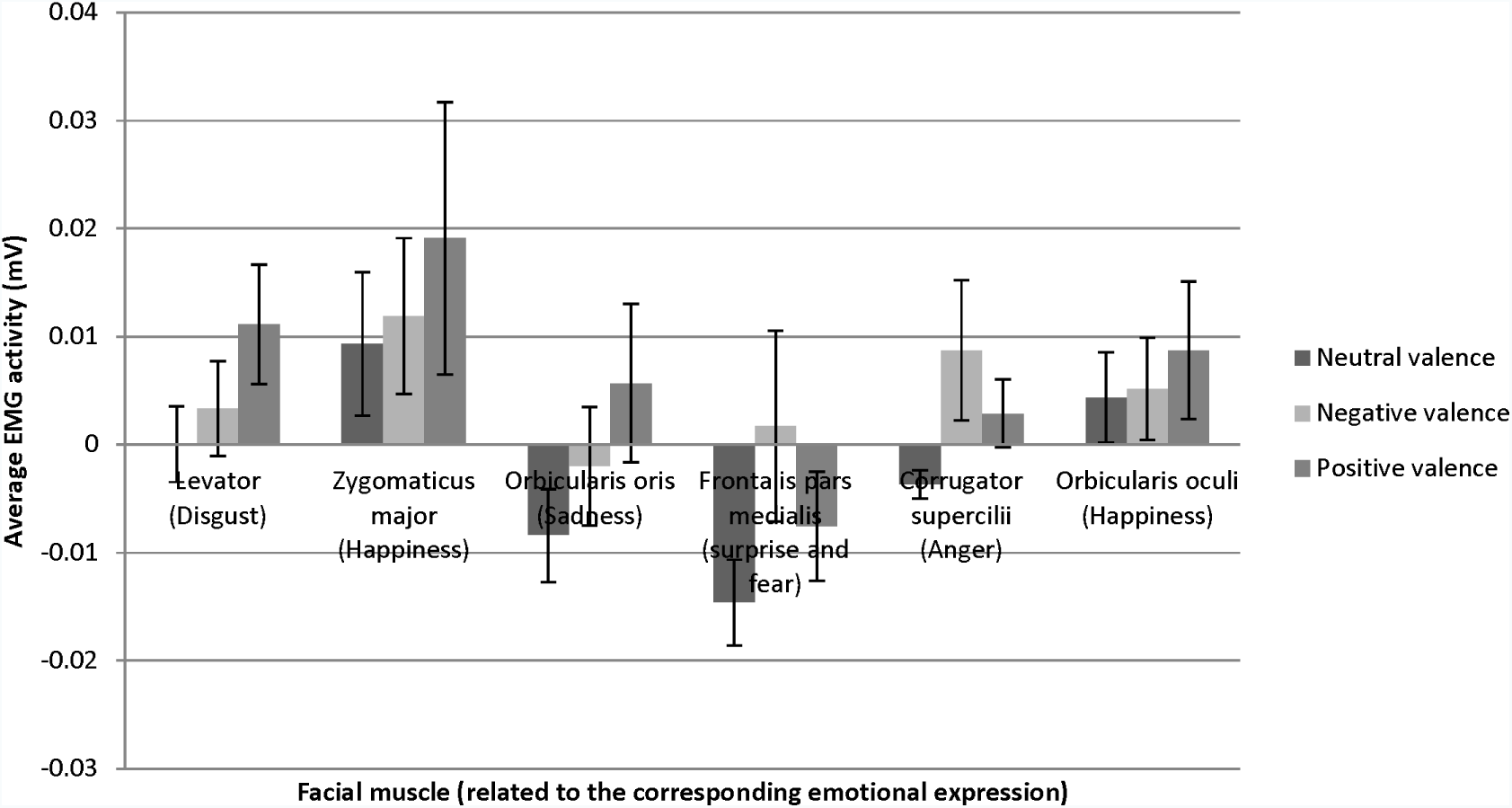
Mean change in EMG activity as a function of emotional expression (on the facial muscle related to this emotional expression) for each social valence of the stimulus.

## Discussion

While we hypothesized that social information will modulate facial mimicry in response to EFE, we found a significant *increase of EMG activity* when EFE were preceded by both positive and negative social label compared to the control situation without social information. The influence of social information on EMG responses was thus *independent* of congruency between the social label and the EFE. Therefore, these findings suggest that the mimicry process is significantly enhanced by the mere presence of social information, independent of its valence.

In respect to the main effect of social information, we supported the hypothesis that social information constitutes a top-down factor that can modulate physiological responses to any EFE. This study is thus in line with previous findings that mimicry of smiles is enhanced in the presence of friends but not in the presence of strangers (Hess et al., 1995) and in the presence of in-group members (Mondillon et al., 2007, Niedenthal et al., 2010; Wang & Hamilton, 2012). In this study, we showed for the first time that social information intrinsically related to the targets of EFE also modulates their processing.

However, we did not show an interaction between social information and EMG responses to EFE, thus failing to replicate the findings in Study 1 and to support the modulatory impact of (in)congruency. Several reasons might explain the absence of interaction between condition and EFE. First, while some studies found an association between subjective and expressive responses to emotional stimuli (Schwartz et al.1980) other failed to find such effect (e.g., Calder et al., 2000; Rottenberg et al., 2002; Sloan et al, 1997). For instance, although spontaneous mimicry leads to greater affective empathy while watching people expressing facial emotions (Stel & Vonk, 2010), Calder and colleagues (2000) showed that, despite their facial paralysis, patients with a Möbius syndrome accurately recognize facial expressions. Therefore, EMG activity after social labels may not be the main determinant of subjective response found in study 1. Second the present findings may suggest that significant increase in EMG activity constitutes a physiological mechanism that allows an enhanced processing of the relevant social information, beyond the object of this information. In other words, the social relevance of individuals has to be processed in a priority manner (relative to its emotional state). This will afterward involve body responses for social nature of the situation, no matter the emotional state of the targets. This hypothesis is supported by findings showing that relative to neutral induction, fear induction before EMG responses increased fear expression when processing fear *and* angry expressions (Moody et al., 2007). Therefore, in line with the latter study, we postulate that EMG responses to EFE are not complete mimicry responses when these EFE are embedded in a socially affective context. One may thus suggest that the EMG activity enhancement after social labels results from the necessity to be alert to any subsequent cues.

### Study 3

Although the social information modulated the subjective (study 1) but not the behavioral responses (study 2) to EFE, study 3 aimed to examine whether rapid neural activity in response to EFE (i.e., P100 and N170 components) is modulated by social context. We hypothesized that the subjective and physiological modulations observed in Studies 1 and 2, respectively, could be related to a modulation of rapid neural components related to visual perception. More precisely, we expected that social information might modulate P100 and N170 components in response to EFE congruent with the social information provided to the participant. Our hypothesis also extends the results reported by Achaibou et al. (2008) that indicated facial mimicry (assessed by EMG activity) is related to P100 and N170 EEG components. Precisely, we expected higher amplitudes of the P100 and N170 in response to congruent trials (e.g. happy face following positive social information) compared to the control trials (no social label). Conversely, we expected lower amplitudes of the P100 and N170 in response to incongruent trials (e.g. happy face following negative social information) compared to the control trials (no social label).

## Method

### Participants

Eighteen healthy subjects (*M*_*age*_ = 21.70; *SD*_*age*_ = 6.64) participated to this study in exchange for course credits. All participants gave written informed consent and had normal or corrected vision and no psychiatric or neurological disorders.

### Stimuli and Procedure

The stimuli were identical to these used in Study 1 and 2. The procedure was also similar except (a) that EEG were recorded during the experiment, (b) that the number of presented blocks was increased and (c) that emotional intensity and valence of the EFE were only rated after the first block for phase 1 (composed of 3 blocks in total) and after the first block for phase 2 (composed of 4 blocks in total). The subsequent blocks for each phase required only passive observation of the EFE (i.e., without subjective responses). Thus, seven blocks were presented to the subjects, including 3 blocks for phase 1 with EFE without social labels (control situations) and 4 blocks for phase 2 with EFE preceded by a social label (either positive or negative). The participants were placed in front of the same screen (CRT 17") as Study 1, but were fitted with the EEG electrodes. EEG recordings were performed during the five blocks of passive exposure to EFE stimuli (from 0 to 2680 msec).

### Data acquisition

Scalp-EEG was amplified using the BIOSEMI Active-Two amplifier and was recorded from 64 electrodes distributed on an elastic cap. The distribution of electrodes was made according to the EEG 10-20 system: the electrodes included Fp1, AF3, AF7, F1, F3, F5, F7, FC1, FC3, FC5, FT7, C1, C3, C5, T7, CP1, CP3, CP5, TP7, P1, P3, P5, P7, P9, PO3, PO7, O1 for the left hemisphere; the equivalent electrodes for the right hemisphere; and FPz, AFz, Fz, FCz, Cz, CPz, Pz, POz, Oz, Iz for the electrodes of the conventional midline sites. Two other electrodes, CMS (Common Mode Sens) and DRL (Driven Right Leg) were respectively used as electrodes of reference and mass (http://www.biosemi.com/faq/cms&drl.htm). CMS was the active electrode while DRL was the passive electrode. We recorded with a sampling rate of 512 Hz and filtered with a low-pass filter of 200 Hz. The signal processing was performed with the software BESA (http://www.besa.de/). The data were all re-referenced to the average reference then filtered with a 1Hz - 40 Hz pass-band filter. For every recording of every subject, a correction of artefacts (eye blinks, etc.) was applied. Artefacts were manually chosen for every subject then automatically corrected for the entire recording. ERPs were averaged for every subject and every condition during the first 1000 ms after the stimulus onset, with a baseline of 200 ms. Then we applied a grand averaging for all the conditions. The component P100 and N170 were studied on the electrodes where their amplitudes were the highest: O1, O2, PO7, PO8, P7, P8, P9, P10.

### Statistical analyses

All the statistical analyses were performed using the STATISTICA 7 software. Mean amplitudes of P100 (time window: 100-140 ms) and N170 (time window: 150-190 ms) were analysed for every condition and for every subject, using repeated-measures ANOVAs and pairwise comparisons. P100 component occurred at the mean latency of 121.90 ms and the mean latency of N170 was 167 ms. Mean amplitudes of both components were analysed as a function of displayed Emotion (7 levels: neutral, anger, sadness, enjoyment, disgust, surprise, and fear), Social Label (3 levels: no social label, positive social label, negative social label) and Hemisphere (2 levels: right, left). Greenhouse-Geisser corrections were applied for variance analysis when violations of sphericity occurred (□ referred to as Greenhouse-Geisser estimate epsilon). The condition of sphericity was verified by a test of Mauchley.

## Results

### P100 component

Analyses revealed a main effect of social label, *F*(1.41, 21.24)= 4.56, *p* < .05, ɛ = .71, indicating that the mean amplitudes of the P100 component differ as a function of the associated social label. EFE preceded by social label, regardless of its valence, elicited larger amplitudes of the P100 compared to EFE presented without social label (but did not differ between positive and negative label). We did not obtain a significant effect regarding hemisphere, *F*(1,15) = .35, ns., nor of the emotion, *F*(6,90) = 1,34, ns. However, the interaction Hemisphere x Emotion was significant, *F*(6,90) = 2,49, *p* < .03, indicating higher amplitudes of P100 on the right hemisphere for the emotions of fear and surprise. No other interaction was significant.

### N170 Component

Similarly to the P100 component, analyses of the N170 component revealed a main effect for social label, *F*(1.34, 20.22) = 6.36, *p* < .05, ɛ = .67. The amplitude was significantly higher for EFE when preceded by positive and negative social label than for EFE without social label (Figure 3) but there was no significant differences between positive and negative social label.

We also observed a main effect for emotion, *F* (6, 90) = 4.61, *p*< .0001, indicating that the N170 amplitudes differed significantly as a function of displayed facial expression toward a higher amplitude for happiness than for other emotions. Analyses showed neither a main effect of the hemisphere, *F*(1,15) = .21, ns, nor any other interaction effect.

**Figure 3.**
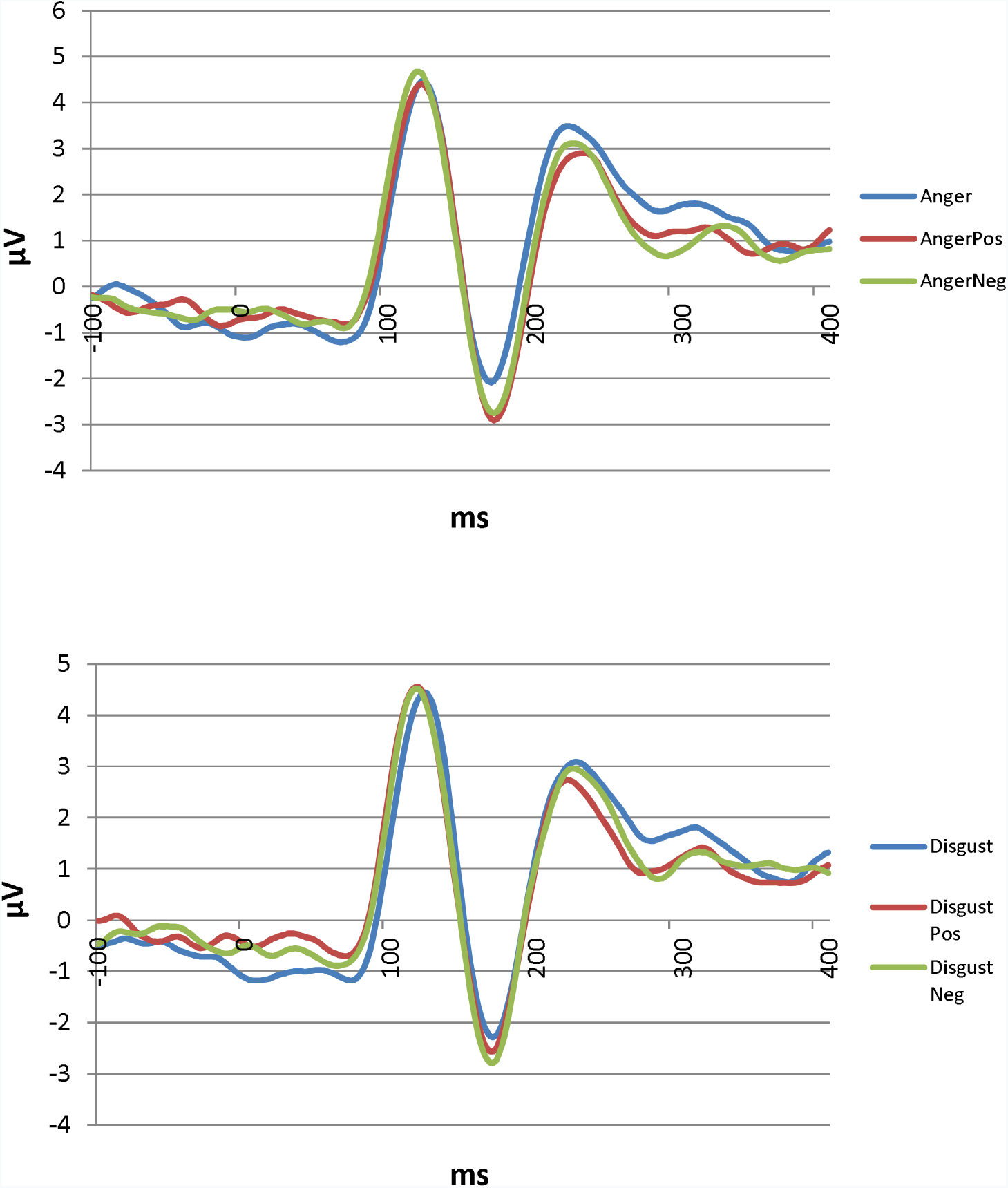
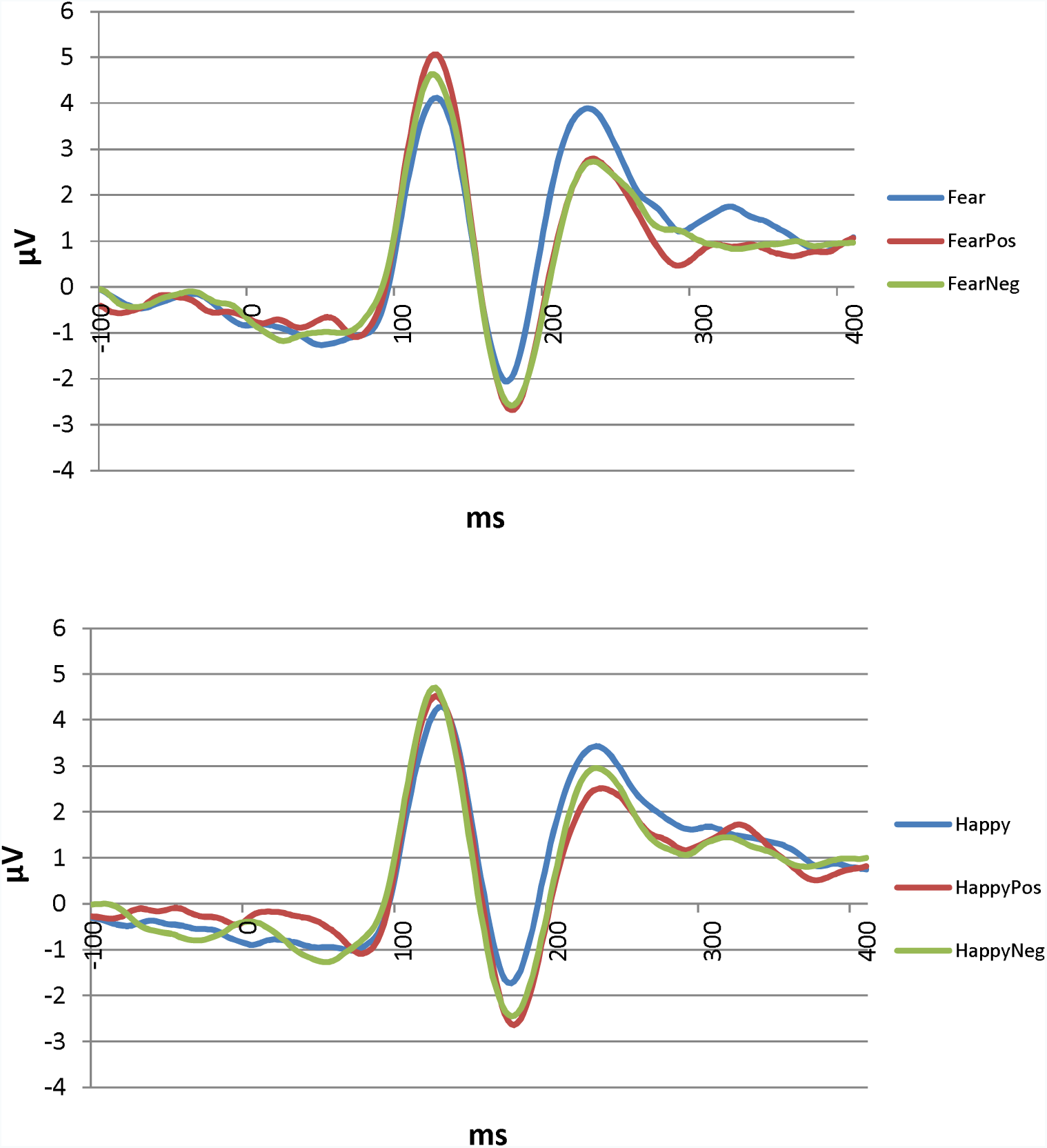
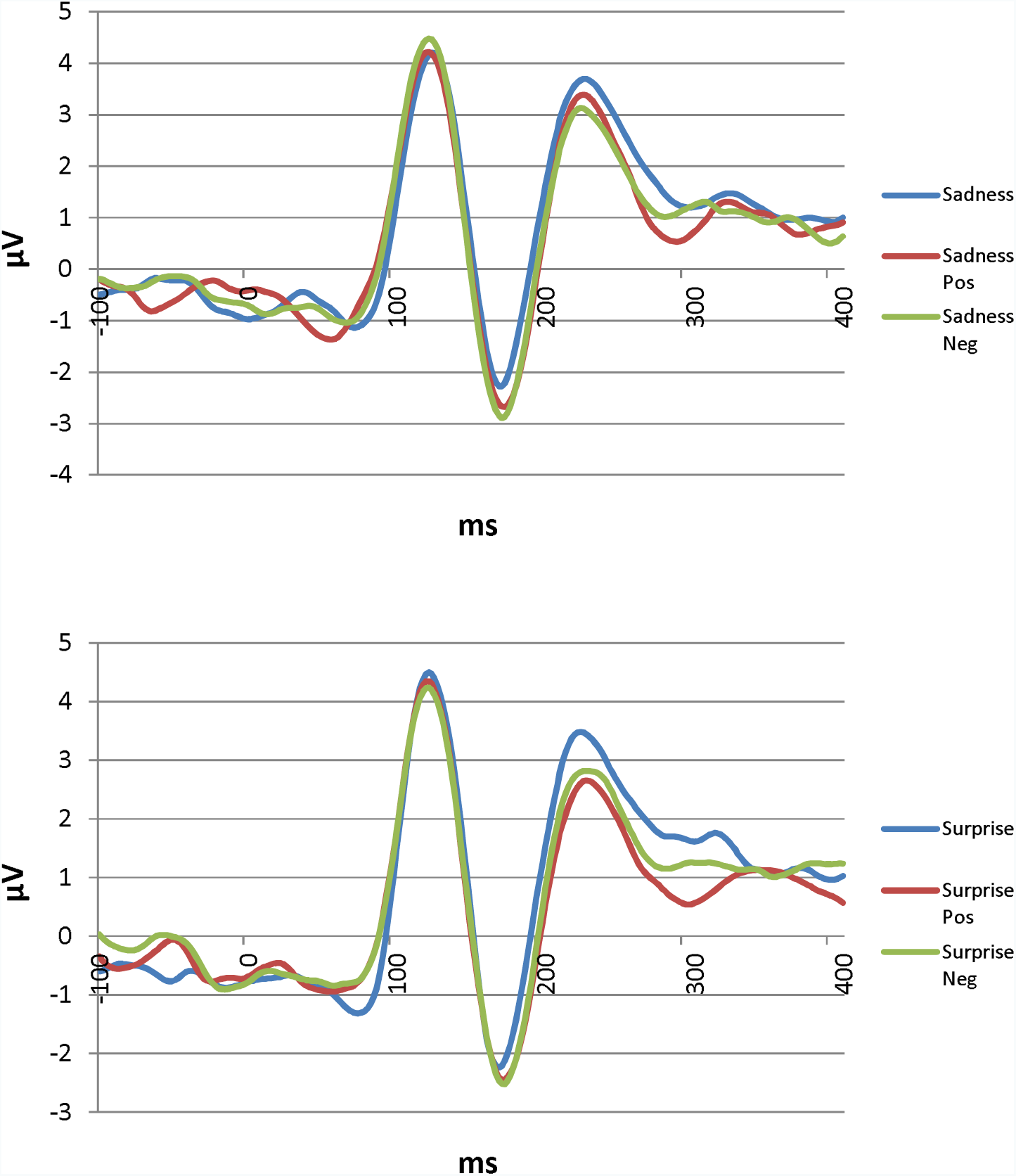
EEG results for P100 and N170 for each emotional expressions preceded or not by positive or negative social valence.

## Discussion

Our results indicate that N170 amplitude was higher for happiness than for other emotions. We also showed higher amplitudes of P100 on the right hemisphere for the emotions of fear and surprise relative to other emotions. More importantly, we expected that early perceptual components could be modulated by (in)congruent social labels for specific EFE (for instance, ERP related to happy faces being modulated by either a positive versus negative social information). We hypothesized higher amplitudes of the P100 and N170 in response to congruent trials compared to control trials (no social label) as well as lower amplitudes of these ERP components in response to incongruent trials compared to control trials (no social label).

Contrary to these hypotheses, there was no significant interaction between social label and EFE. Rather, electrophysiological data are in line with Study 2 and clearly indicate a main effect of social label on both P100 and N170 components. This effect point to a significant increase of both ERP components *irrespective to the social label* associated with any of the EFE, compared to the control situation without social labels. In other words, social information quantitatively increased the neural processes associated with perceptual processing of EFE of the specific individuals but did not qualitatively modulate these processes in regards to the congruency between the social valence and the emotion displayed by the face. Thus, this study is in accordance with the findings of Study 2, which showed a significant increase of EMG activity when EFE were preceded by both positive and negative social label, *irrespective* of the congruency between the social label and the EFE.

### General discussion

The aim of the study was to investigate whether social information about the target expressing a facial emotion could modulate the processing of EFE at subjective, physiological and neural levels. Specifically, we aimed to evaluate the effect of valence congruency between social information about targets and their EFE on the processing of these EFE. The study was based on previous findings showing that perceptual social factors related to the target (e.g., ethnicity) influence the processing of EFE at subjective, physiological and neural levels (e.g., Kubota & Ito, 2007). However, no study had investigated so far the effect of high-level social information about the target on EFE processing by maintaining constant the perceptual features. Finally, no study to date has investigated its effect on embodied facial responses.

In study 1, we supported our hypothesis that social information modulates EFE processing at a subjective level. Indeed, we showed that incongruent valence between social information and EFE reduces the valence and intensity of perceived and felt emotions. Specifically, we showed that expressions of anger, disgust, and fear were perceived as less negative and less intense when preceded by a positive label compared to no label. Conversely, a facial expression of happiness was rated as less positive and less intense when preceded by a negative label compared to no label. We have thus extended upon previous results that suggest social factors related to perceptual features modulate EFE recognition (Mondillon et al., 2007) by showing that high-level social information may impact EFE processing. Therefore, we supported the influence of top-down information (extended to high-order social information) on emotional processing of EFE.

In respect to the results obtained in study 2 at the level of EMG recordings, we supported and extended recent models of embodied cognition assuming that circuits of mimicry are preferentially activated when the observed emotion is socially relevant (Niedenthal et al., 2010; Wang & Hamilton, 2012). More specifically, previous data pointed to a significant modulation of embodiment circuitry by social information such as the presence of friends, but not in the presence of strangers (Hess et al., 1995) or in the presence of in-group compared to out-group members (Mondillon et al., 2007). In the present study, we found weak general EMG activity (including the muscle associated with each specific EFE) when facial expressions were presented alone, without any social label (control condition). However, contrary to our hypotheses, we found a significant increase of EMG activity when expressions were preceded by both positive and negative social label, irrespective of the congruency between the social label and the EFE. This suggests that embodiment circuitries may act as a boost for subsequent cognitive or emotional processes when the situation is socially relevant for the organism, but irrespective of its specific contains (either negative or positive social situations). This supports and extends previous articles suggesting a predominant role of arousal (compared to valence information) during embodiment of emotional cues (Kever, Grynberg, Eeckhout, Mermillod, Fantini, & Vermeulen, 2015). Further studies now have to determine if this physiological variability may have a causal effect on emotional or social behavior (Beffara, Bret, Vermeulen, & Mermillod, 2016).

Moreover, although Study 2 may support the embodiment theory, it remains to be tested whether increased facial reactions after social information priming is accounted for by either affective or social factors. Future study should thus include a neutral social condition in order to better apprehend the respective role of valence and sociality of priming information. Finally, one could argue that EMG activity also results from higher relevance for the self (i.e., involving the observer) during social vs no social priming (Grèzes et al., 2013). Finally, one could also argue that our findings results from cognitive demands to look for facial features that may support or not the social information. Future studies are needed to disentangle these hypotheses.

In Study 3, our goal was to determine if this non-specific boost observed at a psychophysiological level could be related to a modulation, at a neural level, of low level perceptual processes. Our results indicate a main effect of social information as positive and negative social information produced higher amplitudes of P100 and N170 components compared to situations without social information. Therefore, study 3 confirms and extends previous results reported by Achaibou et al. (2008) pointing to a modulation of the P100 and N170 by EFE in relation to EMG activity. Our results are also in line with other findings showing that top-down factors related to the target may modulate neural responses to EFE (Kubota & Ito, 2007; Tortosa, Lupianez, & Ruz, 2013). Finally, it also support fMRI studies have shown that social information modulates the neural correlates of facial expression (e.g., Cloutier, Ambady, Meagher, & Gabrieli, 2012; Singer, Kiebel, Winston, Dolan, & Frith, 2004; Vrtička, Andersson, Sander, & Vuilleumier, 2009; Winston, Strange, O’Doherty, & Dolan, 2002). However, similarly to study 2 and contrary to our initial hypotheses, this effect was independent of the congruency between social information and EFE.

Our findings thus extend the current data by clearly stating that this type of top-down effect cannot be explained by bottom-up perceptual factors (e.g. the perceptual difference between happy and angry faces) since our stimuli were strictly identical at a perceptual level (e.g. an identical smile expressed by a negative versus positive individual). Moreover, our findings revealed that conceptual social information can influence low level perceptual processes of EFE. Therefore, these findings demonstrate that social context has an influence, as early as 120 ms or 170 ms after onset, on low-level perceptual and cognitive processing of emotional expressions.

Although we did not examine the neural activity at a spatial level, we hypothesize that the processing of EFE could be influenced by high-level cortical areas (i.e. orbitofrontal and somatosensory cortices) which would go on to modify perceptual processing of the expression, processed by perceptual areas (Niedenthal et al., 2010). Among neural models that support the influence of top-down factors on perceptual processing (Niedenthal et al., 2010), most of them suggest that perceptual recognition in temporal cortical areas could be influenced by top-down information provided by the orbitofrontal cortex during the recognition of emotions (Barrett & Bar, 2009; Kveraga, Ghuman, & Bar, 2007; Rudrauf et al., 2008). In those models, the early neural activity provided by frontal areas would allow prediction that guides further bottom-up visual processes. Neuroimaging studies with high spatial resolution will have to determine the possible involvement of frontal cortical areas as the origin of the top-down effects reported in the current study.

Moreover, further studies will have to examine whether this modulation of early neural activity could be mediated by a differential allocation of attentional resources to perceptual processes (simple quantitative changes in neural processes) or whether the top-down modulation is able to qualitatively modify the *perception* of emotions. More precisely, our current data does not allow us to determine whether the top-down effects reported here qualitatively modify the perceptual processes occurring at the level of the extrastriate cortex or whether it only constitutes a quantitative boost under the influence of attentional processes. Another important question for further research is to determine the extent to which a non-specific boost observed at the quantitative level of neural and physiological processes (i.e. independently to the congruency between social information and the EFE) is related to further qualitative evaluation observed at a subjective level (and related the congruency between the social valence and the emotion expressed by the target). In other words, it remains to be explained how the interaction between social valence and emotional expressions observed at a subjective level is related to the mere allocation of neural and physiological resources (i.e. irrespective to the social valence of the stimuli) at the early stages of the processing of emotions.

### Conclusion

To conclude, our data provide clear evidence that top-down social information about a target expressing an emotion modifies the perception of this emotion expressed by this target at subjective, physiological and neural levels. In addition, our results suggest that the effect of social label on early neural activity did not systematically vary as a function of its valence (either positive or negative) and the EFE, but, alternatively, produced a global increase of early neural (EEG) and later peripheral (EMG) activity compared to the perception of emotional expressions presented without social context. These findings support and extend recent models of embodiment theory suggesting that the neural and physiological pathways involved in the processing of emotional expressions could be modulated by the social context since the very first stages of perceptual processes (Niedenthal et al., 2010; Wang & Hamilton, 2012). However, further studies should determine whether the effect we obtained with a social label is specific to social information or if other types of relevant and semantic information, emotional or non-emotional, could produce similar top-down influence.

## Acknowledgments

This work was supported by a grant from the Institut Universitaire de France to Martial Mermillod. We thank Pierre Chausse for his help in data acquisition and analysis and Mathias A. Hibbard for English correction of the article.

## Appendix

|  | Positive social label | Negative social label |
| --- | --- | --- |
| Male characters | cheerful sportsman<br>caring father<br>nice nurse | paedophile therapist<br>sadistic killer<br>serial rapist |
| Female characters | loving mother<br>humanitarian doctor<br>passionate teacher | asocial necrophile<br>brutal coach<br>dangerous schizophrenic |

## References

Adolphs, R. (2002). Neural systems for recognizing emotion. Current Opinion in Neurobiology, 12, 169–177.

Achaibou, A., Pourtois, G., Schwartz, S., & Vuilleumier, P. (2008). Simultaneous recording of EEG and facial muscle reactions during spontaneous emotional mimicry. Neuropsychologia, 46, 1104–1113.

Bar, M. (2004). Visual objects in context. Nature Reviews: Neuroscience, 5, 619–629.

Barrett, L. F., & Bar, M. (2009). See it with feeling: affective predictions during object perception. Philosophical Transactions of the Royal Society B: Biological Sciences, 364, 1325–1334.

Barsalou, L. W. (1999). Perceptual symbol systems. Behavioral and Brain Sciences, 22, 577–660.

Beffara B., Bret, A.G., Vermeulen, N., & Mermillod, M. (2016). Resting High Frequency Heart Rate Variability Selectively Predicts Cooperative Behavior. Physiology & Behavior, 164, 417–428.

Beffara, B., Ouellet, M., Vermeulen, N., Basu, A., Morisseau, T., & Mermillod, M. (2012). Enhanced embodied response following ambiguous emotional processing. Cognitive Processing, 13(1), 103–106.

Benson, P. J., & Perrett, D. I. (1993). Extracting prototypical facial images from exemplars. Perception, 22(3), 257–262.

Bublatzky, F., Gerdes, A. B., White, A. J., Riemer, M., & Alpers, G. W. (2014). Social and emotional relevance in face processing: happy faces of future interaction partners enhance the late positive potential. Frontiers in human neuroscience, 8.

Calder, A. J., Keane, J., Cole, J., Campbell, R., & Young, A. W. (2000). Facial expression recognition by people with Mobius syndrome. Cognitive Neuropsychology, 17(1–3), 73–87.

Caharel, S., Courtay, N., Bernard, C., Lalonde, R., & Rebaï, M. (2005). Familiarity and emotional expression influence an early stage of face processing: An electrophysiological study. Brain and cognition, 59, 96–100.

Campanella, S., Falbo, L., Rossignol, M., Grynberg, D., Balconi, M., Verbanck, P., & Maurage, P. (2012). Sex differences on emotional processing are modulated by subclinical levels of alexithymia and depression: a preliminary assessment using event-related potentials. Psychiatry research, 197(1), 145–153.

Clark, V. P., & Hillyard, S. A. (1996). Spatial selective attention affects early extrastriate but not striate components of the visual evoked potential. Journal of Cognitive Neuroscience, 8, 387–402.

Cloutier, J., Ambady, N., Meagher, T., & Gabrieli, J. D. E. (2012). The neural substrates of person perception: Spontaneous use of financial and moral status knowledge. Neuropsychologia, 50(9), 2371–2376.

Dimberg, U. (1990). Facial electromyography and emotional reactions. Psychophysiology, 27, 481–494.

Eimer, M. (2000). The specific N170 component reflects late stages in the structural encoding of faces. NeuroReport, 11, 2319–2324.

Ekman, P., & Davidson, R. J. (1993). Voluntary smiling changes regional brain activity. Psychological Science, 4(5), 342–345.

Ekman, P., & Friesen, W. V. (1976). Pictures of facial affect. Palo Alto, CA: Consulting Psychologists Press.

Fridlund, A.J., & Cacioppo, J.T. (1986). Guidelines for human electromyographic research. Psychophysiology, 23, 567–589.

Grèzes, J., Philip, L., Chadwick, M., Dezecache, G., Soussignan, R., & Conty, L. (2013). Self-relevance appraisal influences facial reactions to emotional body expressions. PloS one, 8(2), e55885.

Harmon-Jones, E., & Peterson, C. (2009). Electroencephalographic methods in social and personality psychology. In E. Harmon-Jones & J. S. Beer (Eds.), Methods in social neuroscience (pp. 170–197). New York, NY: Guilford Press.

Havas, D. A., Glenberg, A. M., Gutowski, K. A., Lucarelli, M. J., & Davidson, R. J. (2010). Cosmetic use of botulinum toxin-A affects processing of emotional language. Psychological Science, 21, 895–900.

Hennenlotter, A., Dresel, C., Castrop, F., Ceballos-Baumann, A. O., Wohlschläger, A. M., & Haslinger, B. (2009). The link between facial feedback and neural activity within central circuitries of emotion—New insights from Botulinum toxin–induced denervation of frown muscles. Cerebral Cortex, 19(3), 537–542.

Hess, U., Banse, R., & Kappas, A. (1995). The intensity of facial expression is determined by underlying affective state and social situation. Journal of Personality and Social Psychology, 69, 280–288.

Hess, U., Adams, R. B., Jr., & Kleck, R.E. (2007). When two do the same, it might not mean the same: The perception of emotional expressions shown by men and women. In U. Hess & P. Philippot (Eds.,) Group dynamics and emotional expression (pp. 33–50). New York: Cambridge University Press.

Hess, U., & Bourgeois, P. (2010). You smile–I smile: Emotion expression in social interaction. Biological Psychology, 84, 514–520.

Kever, A., Grynberg, D., Eeckhout, C., Mermillod, M., Fantini, C., & Vermeulen, N. (2015). The body language: The spontaneous influence of congruent bodily arousal on the awareness of emotional words. Journal of Experimental Psychology: Human Perception and Performance, 41(3), 582–589, doi: 10.1037/xhp0000055

Kveraga, K., Ghuman, A.S., & Bar, M., (2007). Top-down predictions in the cognitive brain. Brain and Cognition, 65, 145–168.

Likowski, K. U., Mühlberger, A., Seibt, B., Pauli, P., & Weyers, P. (2008). Modulation of facial mimicry by attitudes. Journal of Experimental Social Psychology, 44, 1065–1072.

Niedenthal, P. M. (2007). Embodying emotion. Science, 316 (5827), 1002–1005.

Niedenthal, P.M., Mermillod, M., Maringer, M. & Hess, U. (2010). The Simulation of Smiles (SIMS) Model: Embodied Simulation and the Meaning of Facial Expression. Behavioral and Brain Sciences, 33(6), 464–480.

Niedenthal, P. M., Winkielman, P. Mondillon, L., & Vermeulen, N. (2009). Embodiment of Emotional Concepts: Evidence from EMG Measures. Journal of Personality and Social Psychology, 96, 1120–1136.

Mermillod, M., Droit-Volet S., Devaux, D., Schaefer, A., & Vermeulen, N. (2010). Are Coarse Scales Sufficient for Fast Detection of Visual Threat? Psychological Science, 21(10), 1429–1437.

Mondillon, L., Niedenthal, P.M., Gil, S., & Droit-Volet, S. (2007). Imitation of in-group versus out-group members’ facial expressions of anger: A test with a time perception task. Social Neuroscience, 2, 223–237.

Moody, E. J., McIntosh, D. N., Mann, L. J., & Weisser, K. R. (2007). More than mere mimicry? The influence of emotion on rapid facial reactions to faces. Emotion, 7(2), 447.

Murphy, F. C., Nimmo-Smith I., & Lawrence, A. D. (2003). Functional neuroanatomy of emotion: A meta-analysis. Cognitive, Affective, and Behavioral Neuroscience, 3, 207–233.

Pitcher, D., Garrido, L., Walsh, V., & Duchaine, B. C. (2008). Transcranial magnetic stimulation disrupts the perception and embodiment of facial expressions. The Journal of Neuroscience, 28(36), 8929–8933.

Pourtois, G., Dan, E. S., Grandjean, D., Sander, D., Vuilleumier, P. (2005). Enhanced Extrastriate Visual Response to Bandpass Spatial Frequency Filtered Fearful Faces: Time Course and Topographic Evoked-Potentials Mapping. Human Brain Mapping, 26, 65–79.

Pourtois, G., Grandjean, D., Sander, D., & Vuilleumier, P. (2004). Electrophysiological correlates of rapid spatial orienting towards fearful faces.Cerebral cortex, 14(6), 619–633.

Rottenberg, J., Kasch, K. L., Gross, J. J., & Gotlib, I. H. (2002). Sadness and amusement reactivity differentially predict concurrent and prospective functioning in major depressive disorder. Emotion, 2(2), 135.

Rudrauf, D., David, O., Lachaux, J.-P., Kovach, C. K., Martinerie, J., Renault, B., & Damasio, A. (2008). Rapid interactions between the ventral visual stream and emotion-related structures rely on a twopathway architecture. Journal of Neuroscience, 28(11), 2793–2803.

Sadeh, B., Zhdanov, A., Podlipsky, I., Hendler, T., & Yovel, G. (2008). The validity of the face-selective ERP N170 component during simultaneous recording with functional MRI. NeuroImage, 42, 778–786.

Scherer, K. R. (1997). The role of culture in emotion-antecedent appraisal. Journal of Personality and Social Psychology, 73, 902–922.

Schupp, H. T., Flaisch, T., Stockburger, J., & Junghöfer, M. (2006). Emotion and attention: Event-related brain potential studies. Progress in Brain Research, 156, 123–143.

Schwartz, G. E., Brown, S. L., & Ahern, G. L. (1980). Facial muscle patterning and subjective experience during affective imagery: Sex differences. Psychophysiology, 17(1), 75–82.

Schyns, P.G., Goldstone R.L. & Thibaut J.P. (1998). The development of features in object concepts. Behavioral & Brain Sciences, 21(1), 1–54.

Singer, T., Kiebel, S. J., Winston, J. S., Dolan, R. J., & Frith, C. D. (2004). Brain responses to the acquired moral status of faces. Neuron, 41(4), 653–662.

Sloan, D. M., Strauss, M. E., Quirk, S. W., & Sajatovic, M. (1997). Subjective and expressive emotional responses in depression. Journal of affective disorders, 46(2), 135–141.

Smith, M. L., Cottrell, G. W., Gosselin, F., & Schyns, P. G. (2005). Transmitting and decoding facial expressions. Psychological Science, 16(3), 184–189.

Stel, M., & Vonk, R. (2010). Mimicry in social interaction: Benefits for mimickers, mimickees, and their interaction. British Journal of Psychology, 101(2), 311–323.

Tortosa, M. I., Lupiáñez, J., & Ruz, M. (2013). Race, emotion and trust: An ERP study. Brain Research, 1494, 44–55.

Vanlessen, N., Rossi, V., De Raedt, R., & Pourtois, G. (2013). Positive emotion broadens attention focus through decreased position-specific spatial encoding in early visual cortex: Evidence from ERPs. Cognitive, Affective, & Behavioral Neuroscience, 13(1), 60–79.

Vermeulen, N., Mermillod, M., Godefroid, J., & Corneille, O. (2009). Unintended Embodiment of Concepts into Percepts: Sensory Activation Boosts Attention for Same-Modality Concepts in the Attentional Blink Paradigm. Cognition, 112, 467–472.

Vrtička, P., Andersson, F., Sander, D., & Vuilleumier, P. (2009). Memory for friends or foes: the social context of past encounters with faces modulates their subsequent neural traces in the brain. Social neuroscience, 4(5), 384–401.

Wang, Y., & Hamilton, A. F. de C. (2012). Social top-down response modulation (STORM): a model of the control of mimicry in social interaction. Frontiers in Human Neuroscience, 6. doi:10.3389/fnhum.2012.00153

Winston, J. S., Strange, B. A., O’Doherty J., & Dolan, R. J. (2002). Automatic and intentional brain responses during evaluation of trustworthiness of faces. Nature neuroscience, 5(3), 277–283.

